# *Arabidopsis arenosa* influences its microbiome as a serpentine soil adaptation strategy

**DOI:** 10.1101/2025.09.24.678321

**Authors:** Anita Bollmann-Giolai, Michael Giolai, Mellieha Allen, Darren Heavens, Veronika Lipánová, Mark Alston, James Brown, Matthew D. Clark, Paulina Flis, Filip Kolář, Levi Yant, Jacob Malone

## Abstract

There is growing evidence that microbes can facilitate plant growth in metal-rich soils. However, our current understanding of how plants recruit their microbiomes under abiotic stress remains incomplete. Serpentine soils are elementally skewed, with high concentrations of magnesium and nickel, often accompanied by other heavy metals, which can limit calcium availability and present unique challenges to plant growth. These soils are also nutrient-poor, prone to erosion, and have low water-holding capacity. To date, the mechanisms by which plants adapt to serpentine soils remain poorly understood and the role of plant-associated microbiomes in this process has not been described. Here, we focus on *Arabidopsis arenosa* populations adapted to serpentine conditions and investigate their bacterial and fungal microbiomes to better understand the role of plant-associated microbes in serpentine adaptation. We show that serpentine soils harbour distinct plant-associated microbiomes across different plant niches and that the plant genetic background plays a key role in shaping microbial community composition. Finally, we identify serpentine-specific bacterial and fungal variants that may contribute to plant adaptation under these challenging soil conditions.

## Introduction

There is growing evidence suggesting that microbes facilitate plant growth in heavy metal rich soils^1–7^. The plant microbiome influences growth, development and health^8–20^. However, knowledge of how plants attract their microbiome under abiotic stress conditions and the consequences of this for stress survival, is incomplete^9,21,22^. Stress responses such as root morphology or exudation some of can influence plant-microbe interactions^6,21–23^. Plants from highly metal-contaminated sites show distinct microbiomes compared to non-contaminated sites^24,25^. Microorganisms can be used to bioremediate contaminated mining sites^26^. A recent study describes serpentine-derived microbes improving seedling survival in serpentine soils^27^.

Serpentine soils are naturally toxic to plants, globally distributed and derived from ferromagnesium-rich mantle rocks^28,29^. They are the most abundant heavy-metal soils, with elevated nickel, chromium, cobalt and high magnesium, causing low calcium/magnesium ratios and thus magnesium toxicity for plants^28–30^. Serpentine soils often lack essential nutrients^29^, are often rocky, erosion- and drought-prone, with a shallow soil profile with low water-holding capacity^28,29,31^. Such high-stress conditions can have a significant impact on the ecosystem, influencing vegetation composition and leading to a high proportion of endemic species^28,29,31^. Therefore, plants growing on these sites must tolerate both drought and heavy-metals^29,31^.

Serpentine site plants and their adaptation to those conditions have been studied since the 1950s^32–38^. However, in addition to the plants’ inherent capacity to adapt to abiotic and biotic stress, they further rely on their microbiome to enable growth and survival in stressful conditions^8,9,39,40^.

To study plant serpentine adaptation we use an autotetraploid lineage of *A. arenosa*, an outcrosser of the *Arabidopsis* genus, distributed in central and northern Europe as a model plant^33,41,42^. *A. arenosa* repeatedly adapted to serpentine soils and also grows on mining sites^41^. The plant is a facultative excluding metallophyte and also grows on non-contaminated soils^31,43,44^. For serpentine *A. arenosa* populations, genomics data suggests that genes related to the cation transport or chemical homeostasis are under positive selection^33,37^ while a reciprocal transplant experiment documented the fitness advantage of originally serpentine plants in their native substrate^37,38^. Thus, replicated populations, fitness phenotypes and genomic selection signatures make *A. arenosa* to a suitable for serpentine adaptation^45^.

Both environmental and genetic factors influence microbiome formation, but their relative roles under abiotic stress remain unclear. To address this, we examine microbiomes of serpentine and non-serpentine *A. arenosa* populations and assess the relative contributions of environment and plant genome to microbial recruitment We select populations in Austria and the Czech Republic previously characterised for serpentine adaptation^33,37^. Over two seasons, we sampled three serpentine and three nearby non-serpentine sites for microbiomes from different plant niches (i.e., leaf, root, rhizosphere and bulk soil). We also conducted a common garden transplant experiment with all six populations to test genetic effects on microbiome assemblage in both soils. We show that serpentine soils shape microbiomes across niches and that plant genetic background is a major driver of composition. We further identify serpentine-specific bacterial and fungal amplicon sequence variants (ASVs) potentially facilitating adaptation to these challenging soils.

## Results

### Soil ion profiles distinguish serpentine and non-serpentine sites

Serpentine soils are characterized by drought conditions, low nutrient availability, high concentrations of heavy metals and magnesium, and low levels of calcium^46,47^. Magnesium, although being essential to plant growth, has been described to exacerbate the toxicity of other ions, including heavy metals, particularly under low calcium availability^29,48,49^. As a consequence, together with heavy metals, this enhances the toxicity of serpentine soils for plants^50^. Not many plant species grow in such challenging conditions, but *A. arenosa* populations have been previously described to show local adaptation to serpentine soils^37,38,45^ (**Fig. 1A**).

**Fig. 1.**
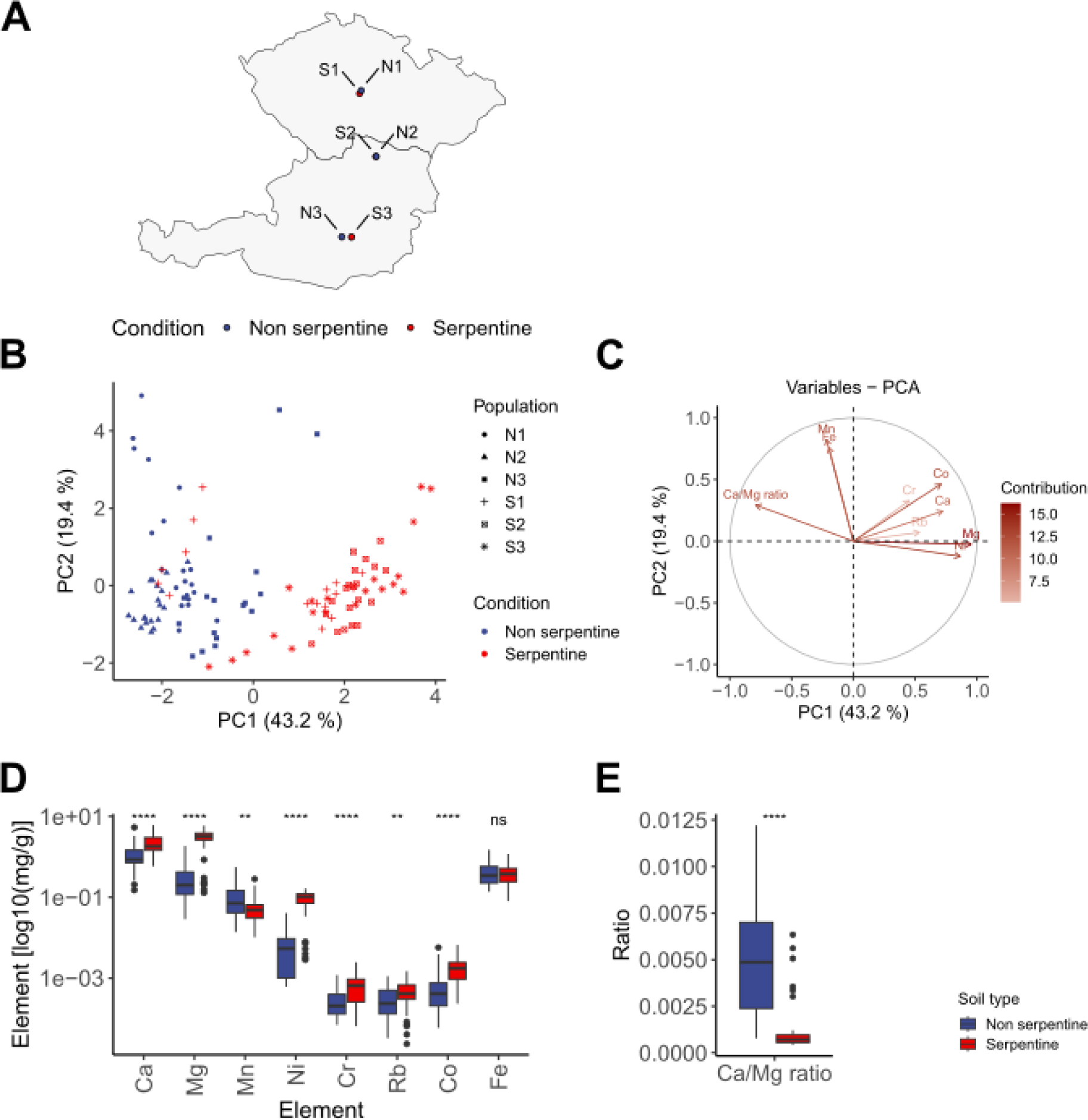
Serpentine soils, and their associated *A. arenosa* populations, display elevated metal ion concentrations and a lower calcium to magnesium ratio than non-serpentine samples. **(A**) Geographical locations of serpentine and non-serpentine populations in Austria and the Czech Republic. **(B)** PCA of soil ion composition showing clear separation of serpentine and non-serpentine sites. **(C)** PCA biplot illustrating contributions of individual ions and the Ca/Mg ratio to the first two principal components. **(D)** Ion concentrations obtained from soil profiling. Labels indicate statistically significant differences between serpentine and non-serpentine soils (t-test). **(E)** Difference in Ca/Mg ratio between serpentine and non-serpentine soils (t-test). (*p* < 0.0001****, *p* < 0.001***, *p* < 0.01**, *p* < 0.05*; not significant: ns)

We selected three pairs of populations established in either serpentine or non-serpentine soils, which are growing in spatial proximity in Austria and the Czech Republic (**Fig. 1A**). To confirm the soil ion profiles of the selected sites we collected soil material along with microbiome sampling and analysed the ion composition data of all sites (**Fig. 1B-E, Extended Data File 1**). Principal component analysis (PCA) of soil ion composition showed that that serpentine and non-serpentine soils separated well along PC1, which explained 43.2 % of variance (**Fig. 1B**). Magnesium, calcium and rubidium as well as the heavy metals nickel, chromium and cobalt contributed most to the separation (**Fig. 1C**). The magnesium concentration showed the strongest association with serpentine soils along PC1. We also analysed the Ca/Mg ratio and found that the ratio was consistently lower in serpentine than non-serpentine soils (**Fig. 1C-E**, **Extended Data File 1**). This pattern agrees with previously described characteristics of serpentine soils^32^.

### Microbiome community structure across serpentine and non-serpentine niches

We collected bulk soil (adjacent to plants), rhizosphere, root and leaf material in spring and fall 2018 from five plants per population. We constructed and sequenced 16S and ITS Illumina amplicon libraries and assigned ASVs^23,51^, removing ASVs below a total relative abundance of 0.1%. We performed a redundancy analysis (RDA) to assess community structure across all niches using genus-level taxonomic resolution (**Fig. 2, Extended Data File 2**). Here, we observed stronger niche differentiation in bacterial communities (**Fig. 2A, Extended Data File 2**) compared to fungal communities (**Fig. 2C, Extended Data File 2**). In both bacteria and fungi, leaf-associated communities were clearly distinct from those in bulk soil, rhizosphere, and roots (**Extended Data File 2**).

**Fig. 2.**
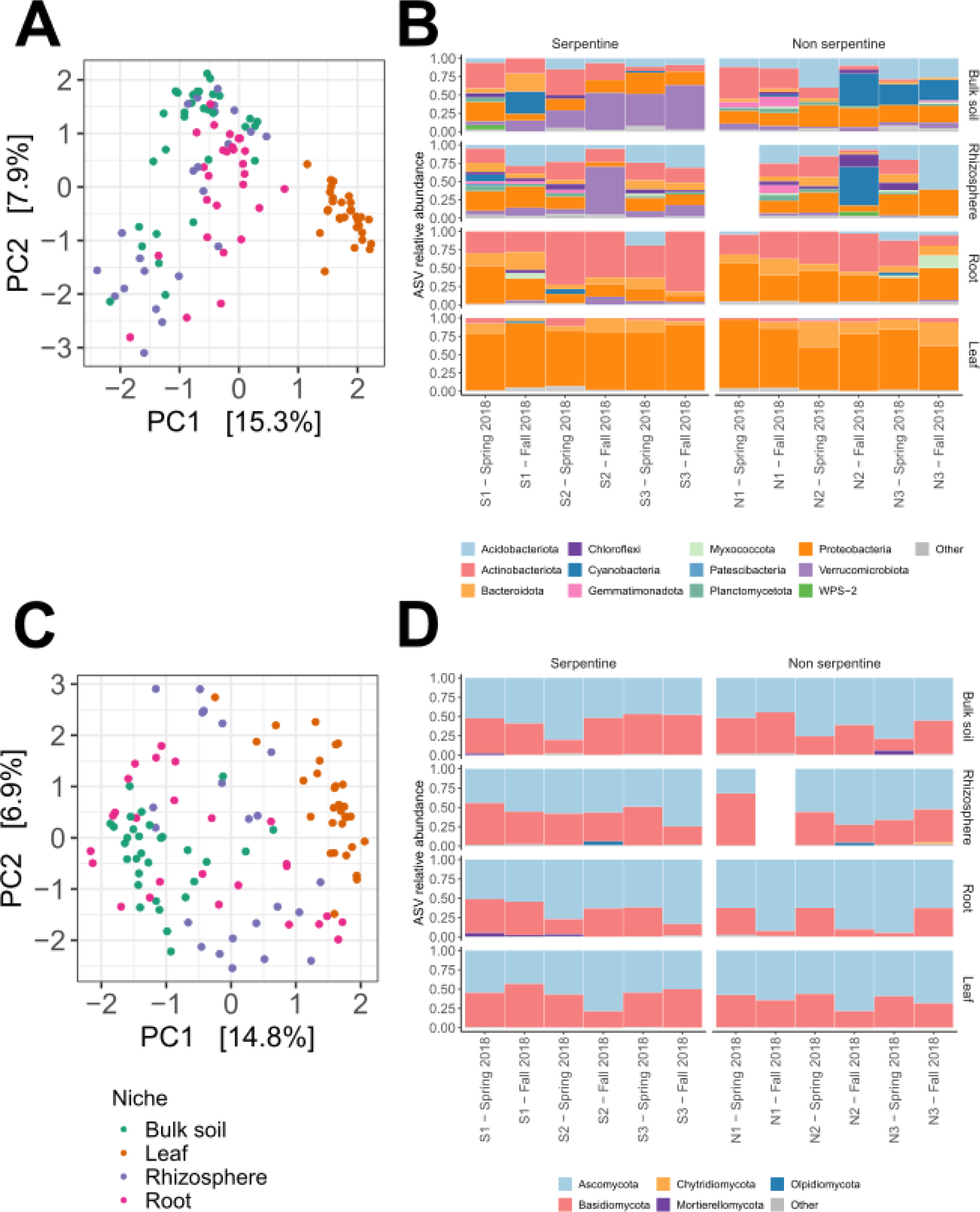
Higher bacterial variation in below-ground niches and higher uniformity for fungal communities across all niches. **(A)** RDA of bacterial communities in bulk soil, rhizosphere, root, and leaf niches. **(B)** Relative abundance of bacterial phyla across all populations and seasons (spring and fall 2018). **(C)** RDA of fungal communities in bulk soil, rhizosphere, root, and leaf niches. **(D)** Relative abundance of fungal phyla across all populations and seasons (spring and fall 2018). Fall 2018 was used as representative season for A and C. (Phyla with a total relative abundance of less than 2 % were grouped as ‘Other’.)

ASVs were grouped by phylum and illustrated in bar plots to understand bacterial and fungal microbiome composition across populations and seasons (**Fig. 2B,D**). Bar plots indicated that the phylum-level composition of bacterial and fungal communities was largely similar between serpentine and non-serpentine sites, but showed niche specific differences between bacteria and fungi. Multiple bacterial phyla were present across all four niches, whereas fungal communities were dominated by few highly abundant phyla. This aligns with previous findings that bacterial niches exhibit greater diversity and niche differentiation compared to fungi^52^. For bacteria, 11 and 13 phyla were detected in bulk soil and rhizosphere compared to 8 in roots and 5 in leaves (**Fig. 2B**). In the bulk soil compartment Cyanobacteria, Proteobacteria, Actinobacteria, Verrumicrobiota and Acidobacteriota shared equal relative abundances (22.71 % - 16.12 % of assigned ASVs respectively). The same pattern was observed for the rhizosphere, with some changes in abundances and an increase of Cyanobacteria from 22.71 % to 31.35 % of assigned ASVs. Roots were dominated by Actinobacteria (44.19 %) and Proteobacteria (31.59 %), whereas leaves were highly enriched in Proteobacteria with 78.22 % of assigned ASVs (**Extended Data File 3**). This indicates potential niche specific shifts and composition of bacterial taxa from bulk soil to the rhizosphere, as well as roots to leaves. Fungal communities exhibited less clear niche differences (**Fig. 2D**). Bulk soil and rhizosphere contained 3 and 4 phyla while roots and leaves contained 3 and 2 phyla respectively. Ascomycota and Basidiomycota together accounted for the majority of relative abundances across all niches. Ascomycota enriched for an average percentage of 58.74 %, 56.16 %, 71.37 % and 60.26 % from bulk soil, rhizosphere, roots to leaves. For Basidiomycota these loads were 39.67 %, 41.55 %, 27.2 % and 39.24 % respectively (**Extended Data File 3**). The observation of differences in microbial composition between niches of serpentine and non-serpentine plants agrees with previous findings by Botha *et al.*^53^.

### Differential abundance analysis reveals serpentine-specific enrichment of microbial taxa

We hypothesised that functional effects of a potential microbiome facilitated serpentine adaptation likely occur at a finer taxonomic resolution than the phylum level (e.g., genus or species)^53,54^.

Therefore, we tested whether individual ASVs influenced differences in community composition by examining their occurrence across plants and populations: ASVs present in all populations were considered conserved, those found in two of three populations were classed as shared, and those restricted to a single population as unique (**Fig. 3A**). Analyses were performed for each niche, for serpentine and non-serpentine sites, and for bacterial and fungal communities. Across 14,021 bacterial and 5,672 fungal ASVs we found that most bacterial and fungal ASVs were unique to individual populations (**Fig. 3A, Extended Data File 4**). A fraction of 1.10 % – 9.17 % of serpentine bacterial ASVs and 7.49 % - 9.51 % for serpentine fungal ASVs was conserved across niches. For non-serpentine niches these numbers were lower with 0.31 % - 7.48 % and 3.16 % - 8.52 % for bacterial and fungal ASVs across niches respectively. We observed the highest number of conserved ASVs for fungal serpentine microbiomes (**Fig. 3A, Extended Data File 4**). Despite the large number of ASVs that were unique to individual populations, the results indicate that plants growing on serpentines harbour a conserved subset within their microbial communities that potentially contributes to serpentine adaptation^24,25,53^.

**Fig. 3.**
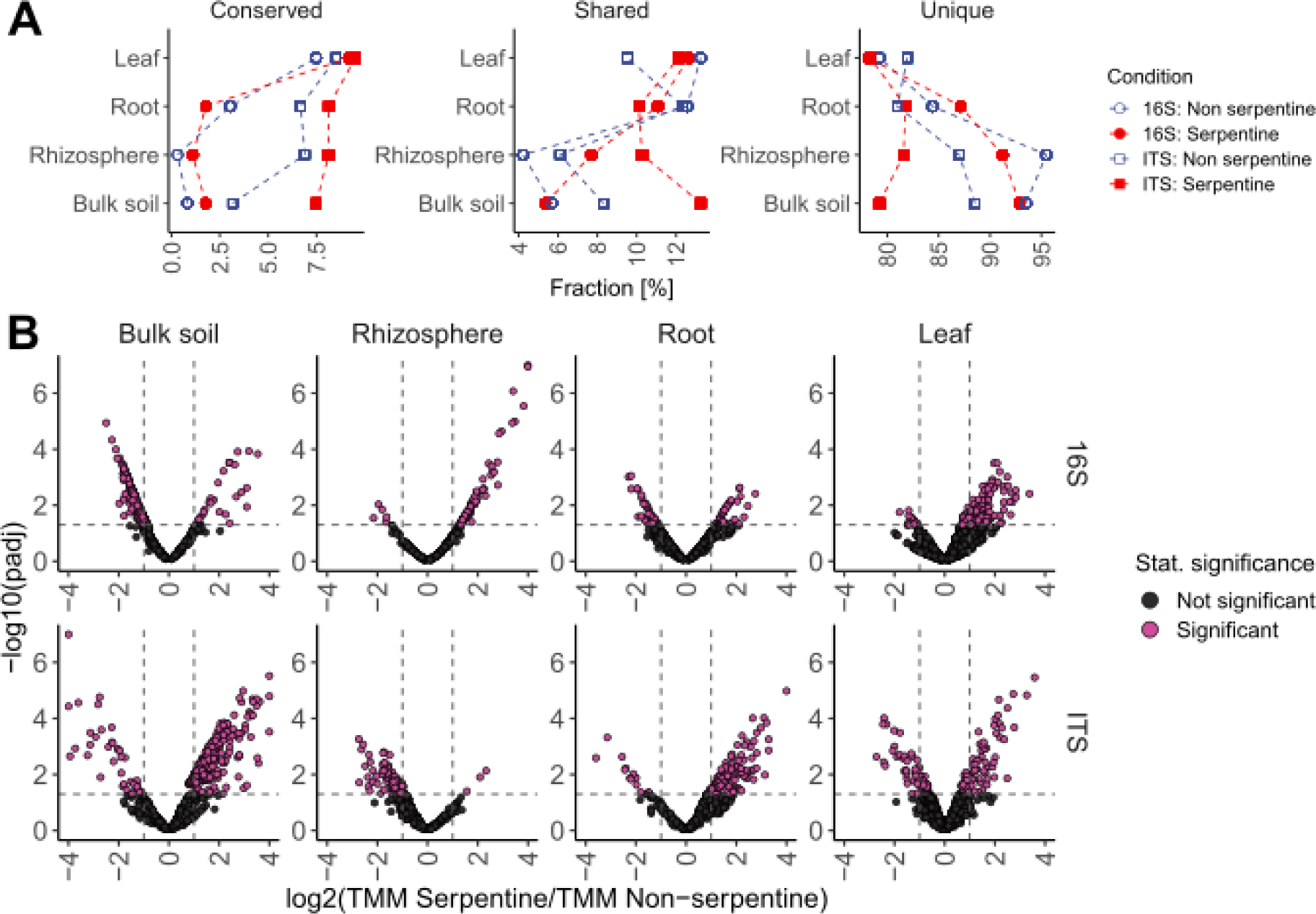
Similarity of bacterial and fungal microbiomes across plant populations based on ASV presence-absence and differential enrichment. **(A)** Distribution of bacterial and fungal ASVs across niches, categorised as conserved (present in all three populations), shared (present in two populations), or unique (restricted to a single population). Fungal ASVs were more conserved than bacterial ASVs under both serpentine and non-serpentine conditions. **(B)** Differential abundance of conserved bacterial and fungal ASVs between serpentine and non-serpentine soils. Volcano plots show significantly enriched ASVs.

To test whether these patterns reflect statistically significant compositional shifts between serpentines and non-serpentines, we performed differential abundance analysis using the microbiomeMarker package^55^ (**Fig. 3B**). A total of 13,895 bacterial and 5,618 fungal ASVs met prevalence criteria^55^. Of these we found that 13.74 % of bacterial and 22.59 % of fungal ASVs were statistically significantly enriched (**Extended Data File 5, Extended Data File 6**). Whereas 2.84 % of 13.74 % of bacterial ASVs across all niches enriched for serpentines. For the fungi this pattern was the other way around and the majority of ASVs enriched for serpentines with 16.02 % of 22.59 %. These results indicate that fine-scale taxonomic shifts accompany the previously observed broader community differences. We next analysed the genera to which serpentine-enriched ASVs belonged. For each genus, we counted the number of ASVs and calculated the log_2_-fold-ratio of serpentine to non-serpentine ASVs. (**Extended Data File 5, Extended Data File 6**). For bacteria in bulk soil, no genus showed a substantially higher number of ASVs in serpentine sites. In the serpentine rhizosphere, *Candidatus Udaeobacter* was the most enriched genus with 47 ASVs compared to none in non-serpentine sites, followed by *Vicinamibacterales* with 16 vs. 0 ASVs. In roots, genera generally had lower ASV numbers, with *Streptomyces* showing 4 vs. 0 ASVs compared to serpentine sites. In leaves, *Hymenobacter* and *Methylobacterium–Methylorubrum* counted more ASVs for serpentines, with 45 vs. 0 and 25 versus 0 ASVs to non-serpentines, respectively. For fungi, *Mytilinidion* (19 vs. 0), *Knufia* and *Russula* (both 13 vs. 0) were most abundant in bulk soil, while no genera were notably enriched in the rhizosphere. In roots, *Penicillium* (7 vs. 0), *Knufia*, *Sebacina,* and *Tomentella* (all 4 vs. 0) were enriched, and in leaves, *Fellozyma* was observed with 4 vs. 0 ASVs compared to non-serpentines (**Extended Data File 7**). Consolidating known literature about these genera we found commonalities among bacteria and fungi that potentially contribute to microbiome mediated serpentine adaptation. I.e., *Streptomyces* and *Methylobacterium–Methylorubrum* bacterial genera have been described to be beneficial for plant performance and growth^56–58^. *Sebacinales* fungi are potential plant growth promoting endophytes^59^. *Penicillium* fungi are described as decomposers of organic materials^60^, and *Tomentella* and *Russula* have been recognised as ectomycorrhizal fungi that are important in nutrient cycling^61,62^. Hence, bacterial and fungal genera could together contribute to serpentine adaptation by facilitating nutrient availability and promoting plant growth in serpentines. This agrees with previous findings where serpentine plants have been described to associate with arbuscular mycorrhizal assemblages^63^.

### Serpentine microbiomes are better retained by native plants under reverse-transplant conditions

Previous studies have shown that plant genotypes can influence the composition of their associated microbiomes^64–67^. However, environmental factors also significantly impact microbial community composition and environmental stressors can influence the recruitment of specific microbial taxa that may aid in plant adaptation^68^. It remains unclear to what extent plants actively recruit microbes or whether observed effects are environmentally driven.

To examine potential recruitment effects, we predicted that non-serpentine genotypes, which are not typically adapted to serpentine soils^38^, would exhibit different responses compared to serpentine genotypes under serpentine conditions. To test this, we performed a reciprocal transplant experiment in which serpentine and non-serpentine populations were grown in both serpentine and non-serpentine soils. As in the field experiment, we collected five plants for each condition, performed 16S and ITS Illumina sequencing and called ASVs^69^. We then compared ASV abundances of serpentine and non-serpentine populations grown in their native soil type with those in the transplanted (i.e., non-native) soil using Bray-Curtis similarity (**Fig. 4A,B**). We observed that microbiomes between native and non-native conditions differed substantially, yet not completely, between populations. Fungal microbiomes were more similar than bacterial microbiomes in the bulk soil and the root compartments (**Fig. 4C, Extended Data File 8**). To test whether serpentine and non-serpentine plants recruit similar microbiomes under serpentine conditions, we compared the log_2_ fold-changes of ASVs observed in the field with those measured in the transplant experiment, focusing on serpentine-to-non-serpentine enrichment. Log_2_ fold-change values were compared using Spearman’s rank correlation coefficients (**Fig. 4C**). The analyses showed that log_2_ fold-changes in serpentine plants from the transplant experiment agreed more closely with field data and preserved the patterns observed in the field better than those of non-serpentine plants. For serpentine plants, Spearman correlation coefficients with the field data were 0.71, 0.66, and 0.59 for bulk soil, rhizosphere, and root, respectively (all statistically significant: *p* ≤ 2×10^-16^, *p* = 5×10^-3^, and *p* = 5×10^-4^). For non-serpentine plants, correlations were weaker with 0.62 and 0.46 for bulk soil and root (bulk soil *p* = 1.8×10^-12^, root *p* = 1.9×10^-2^). These results indicate that serpentine plants maintained a microbiome that more closely resembles the field serpentine gradient than non-serpentine plants. No statistically significant correlations were observed in leaves, which potentially reflects aboveground effects of growth under controlled conditions rather than patterns present in the field.

**Fig. 4.**
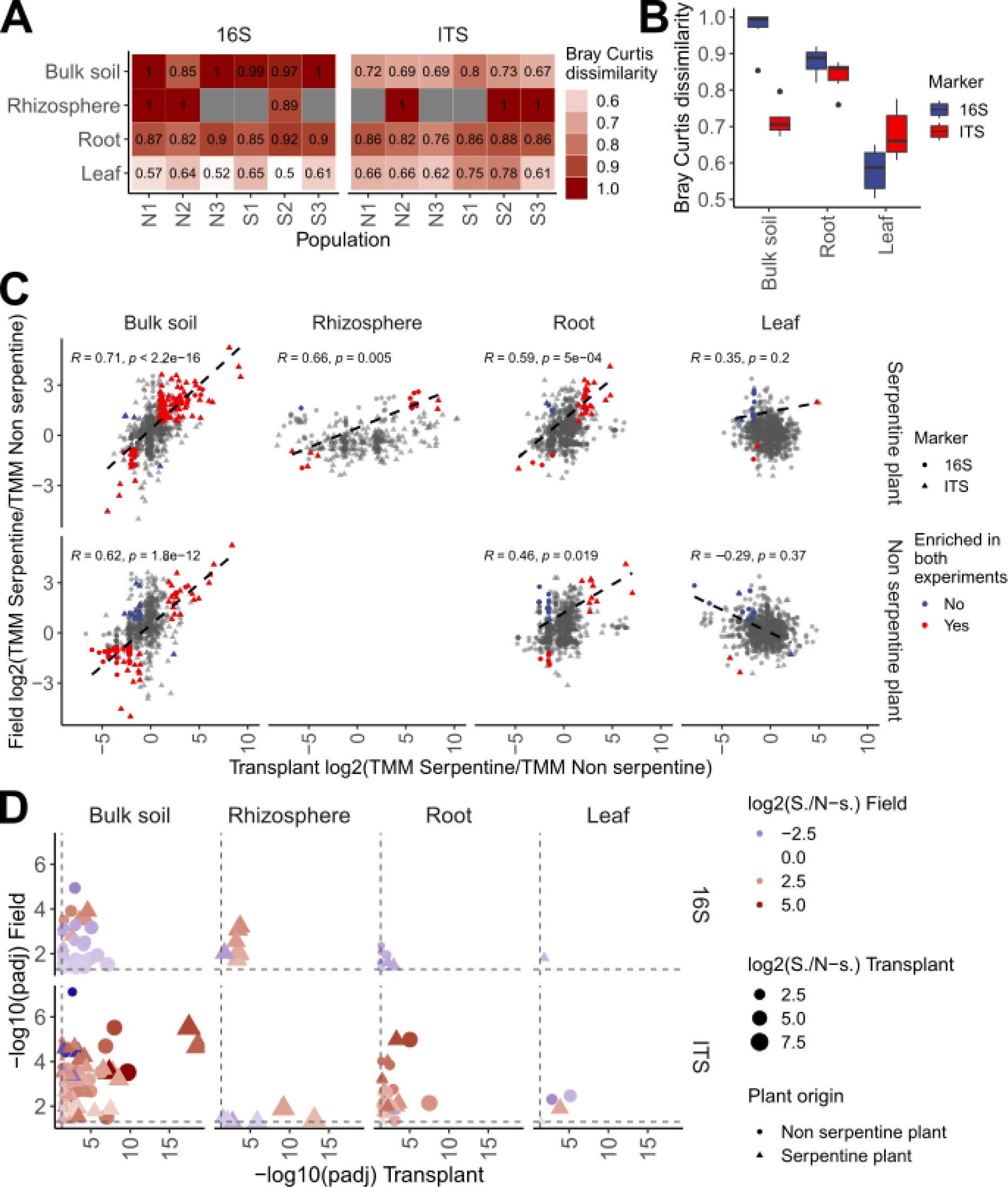
Microbiome responses in a reciprocal transplant experiment. **(A,B)** Bray-Curtis dissimilarity shows that microbiomes of transplanted plants differ from, but partially retain, those in native soils, with fungi showing greater similarity across compartments than bacteria. The heatmap in **(A)** shows Bray-Curtis dissimilarity scores per population and niche for bacteria and fungi, boxplot **(B)** visualises the Bray-Curtis dissimilarity scores next to one another across niches. **(C)** Spearman correlations of log2 fold-changes reveal that serpentine plants more closely mirror field enrichment patterns than non-serpentine plants, particularly belowground. **(D)** Differential abundance analysis identifies ASVs enriched in serpentine versus non-serpentine soils, with overlap to field-enriched ASVs highlighting candidates potentially contributing to serpentine adaptation.

We hypothesised that ASVs with a potential role in facilitating serpentine adaptation would be statistically significantly serpentine-enriched in the field and the transplant data. We therefore compared statistically significantly enriched bacterial and fungal ASVs from the field with those from the transplant experiment (**Fig. 3D**). Overall, serpentine plants showed stronger overlap with field data than non-serpentine plants. For bacteria, only three ASVs in bulk soil were serpentine-enriched in populations of non-serpentine origin, whereas populations of serpentine origin showed greater overlap, with six ASVs in bulk soil and nine in the rhizosphere. For fungi, the numbers were generally higher. In serpentine-origin plants, we observed 90 overlapping ASVs in bulk soil, one in leaf, two in the rhizosphere, and 24 in root. In contrast, non-serpentine-origin plants showed fewer overlaps, with 21 in bulk soil, none in leaf or rhizosphere, and 10 in root.

We found that fewer bacterial ASVs were enriched under serpentine conditions, whereas fungi showed a greater number of consistently enriched ASVs, both in the field and in serpentine plants during the transplant experiment. This overlap highlights fungal ASVs as potential contributors to serpentine adaptation, particularly in belowground compartments such as bulk soil, rhizosphere, and root. Among bacteria, only a single genus, *Candidatus Udaeobacter*, was enriched in both the rhizosphere and bulk soil. In contrast, we identified 73 enriched associations with fungal taxa, 55 of which could be assigned at the genus level (the remainder assigned at higher taxonomic ranks such as phylum, order, or class). We also quantified the number of ASVs per genus and, consistent with the field data, observed overlap with genera already detected in the natural serpentine community (**Extended Data File 9**). For example, we identified six ASVs in *Penicillium*, three in *Russula*, and one each in *Mytilinidion* and *Tomentella*. Consolidating literature further we also detected the fungal genus *Exophiala*, which has been described to promote plant growth under nickel stress, a major separator of serpentine conditions in our PCA^70^ We also found the fungal genus *Phialocephala*, which is a plant growth promoter by enabling nutrient availability^71^. Previous work has shown that serpentine plants associate with distinct arbuscular mycorrhizal assemblages that may help mitigate metal stress^63^. Our findings therefore indicate potential new fungal partners involved in serpentine stress adaptation. Moreover, fungi associated with serpentine plants are often highly diverse and may include specialized taxa that confer advantages to plants growing under serpentine conditions^72,73^.

### Plant genotype is a stronger factor shaping microbial communities, compared to the environment

The extent to which plant-associated microbiomes are actively recruited by the host, shaped by environmental conditions, or both remains unclear^74^. Considering enrichment patterns and population differences between our field and transplant experiment, we concluded that microbiome assembly results from a complex interplay between plant genotype and environmental factors, with both likely contributing^75,76^. Disentangling the relative role of plant host and environment is essential to determine whether microbiome-mediated stress adaptation is genetically encoded and thus potentially subject to evolution or primarily environmentally induced.

To disentangle the effects of host genotype and environment on microbiome recruitment, we focused on the statistically significant, differentially enriched ASVs identified in the field experiment. Genotype data from populations revealed clear population separation in MDS analyses of k-mers from both individual genome sequencing^37^ (**Extended Data File 2**). We then applied generalized linear mixed models (GLMMs) to test whether ASV abundances were explained by environment (serpentine vs. non-serpentine), genotype (population), or their interaction. Model contributions were quantified using R^2^ values. Across all four niches, the combined effect of genotype and environment accounted for the largest proportion of variance. Importantly, for both bacterial and fungal communities, genotype-only models consistently explained more variance than environment-only models. These results indicate that microbiome assembly is shaped by both host genotype and environment, with a stronger contribution from genotype (**Fig. 5**). This pattern supports the idea that serpentine adaptation reflects a long-term evolutionary process involving both plants and their associated microbiomes.

**Fig. 5.**
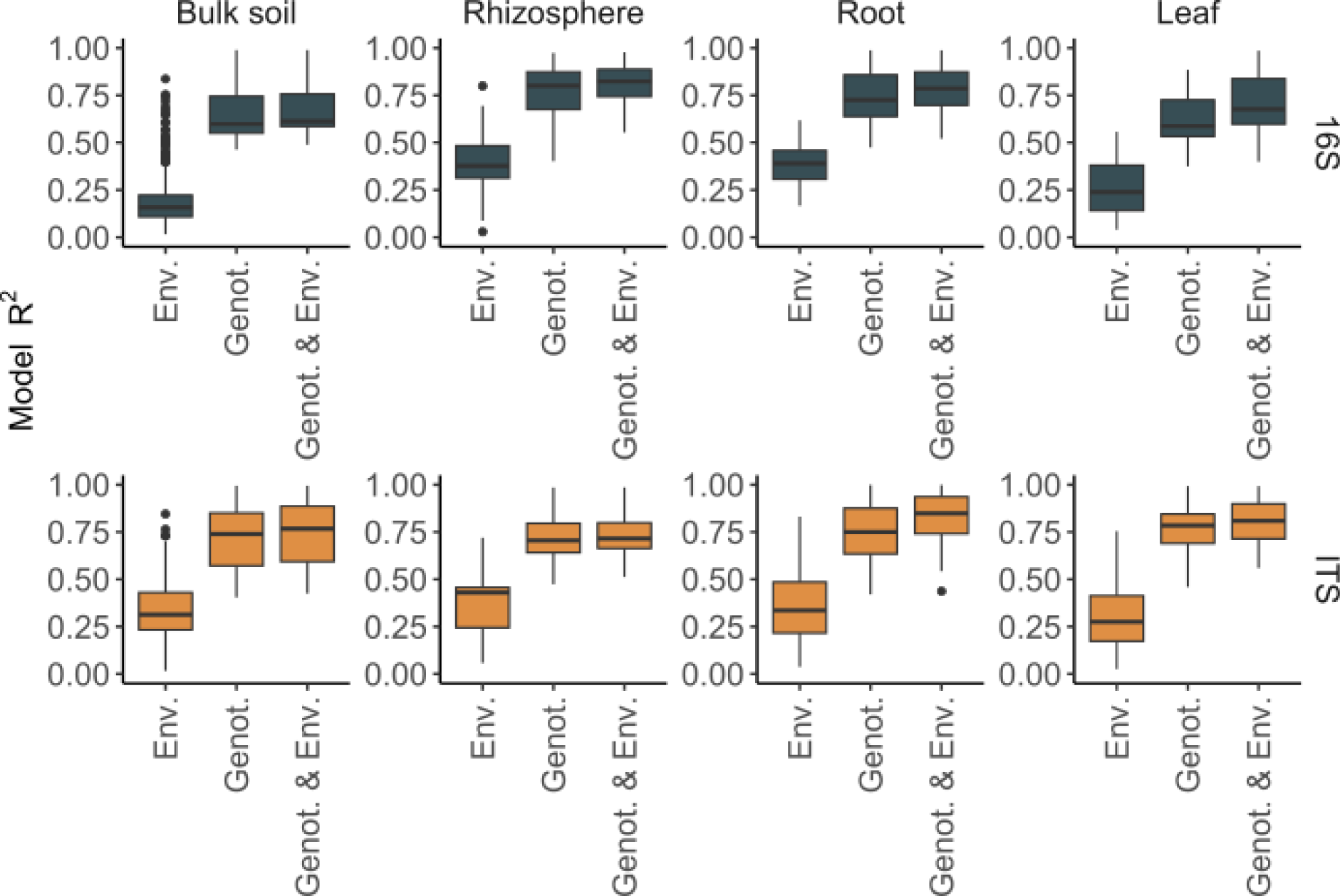
Contributions of genotype and environment to microbiome composition. Explained variance (R^2^) of ASV abundance is shown for statistically significant models predicting ASV relative abundance, with genotype and environment as fixed factors and time as a random factor. While the combination of genotype and environment best predicts ASV abundance, genetic effects consistently exceed environmental effects in shaping microbiome assembly.

## Discussion

In this study, we conducted an in-depth examination of the microbiome communities associated with *A. arenosa* plants growing in the challenging environment of serpentine soils. Serpentine soils impose multiple simultaneous stresses, including low calcium-to-magnesium ratios, high heavy metal loads, and nutrient deficiencies^32^. We show that serpentine soils differ from non-serpentine soils in their ionomic composition, most notably by a lower calcium-to-magnesium ratio. We further demonstrate that microbiomes of serpentine and non-serpentine plant populations differ in both bacterial and fungal composition across all tested compartments, from bulk soil to rhizosphere, roots, and leaves. In a reciprocal transplant experiment, exposing plants of serpentine and non-serpentine origin to both their native and non-native soils, we observe distinct microbial compositions depending on plant origin and soil type.

By carefully confirming field observations with a transplant experiment, we identify several ASVs and genera that can contribute to microbiome-mediated serpentine adaptation. Notably, many of the genera that we find as statistically significantly associated with serpentine soils are described to enhance either nutrient availability or to promote plant growth. This involves the coordinated contribution of both bacteria and fungi. Beneficial bacterial genera that we identify, such as *Streptomyces* and *Methylobacterium–Methylorubrum* enhance plant growth^56–58^, while fungal taxa including *Sebacinales*, *Penicillium*, *Tomentella*, and *Russula* support nutrient cycling, organic matter decomposition, and soil nutrient acquisition^59–62^. Additionally, *Exophiala* and *Phialocephala* promote plant growth under nickel stress and improve nutrient availability^70,71^. Together, these microbes likely facilitate plant survival and growth under serpentine stress, complementing known associations with arbuscular mycorrhizal fungi^63^. Considering that serpentine soils are nutrient-poor and subject to heavy metal stress^32,46,47^, the coordinated presence of microbial taxa that enhance nutrient availability, promote plant growth, and mitigate metal stress is likely crucial for plant survival under these conditions^9^. From an ecological and evolutionary perspective, these results are particularly impactful because they demonstrate a potential mechanism by which plants can extend their adaptive capacity through microbial partnerships.

In previous findings Konečná *et al.*^37^ reported that *A. arenosa* exhibits an adaptive phenotype in serpentine soils, with this adaptation linked to 60 candidate genes, notably including the calcium channel *TPC1*. Whereas *TPC1* has not been connected to microbiome recruitment in detail^37^, calcium channels play a role in plant signaling, microbe interactions^77^. In addition to confirming the importance of plant genetics in serpentine adaptation, we further show that the host genotype exerts a stronger effect than soil environment on the microbiome composition. This suggests that heritable plant traits influence microbial recruitment in ways that could reinforce adaptive phenotypes under stressful conditions. Such findings provide direct evidence supporting the holobiont concept, in which the plant and its microbiome function as an integrated adaptive unit^39^. Recognising the plant and its microbiome as an integrated adaptive unit opens new avenues for research and provides a roadmap for sustainable strategies to enhance plant health and productivity in challenging ecosystems^74,76,78,79^.

In conclusion, our work highlights that microbial contributions to plant adaptation are both specific and compartmentalized. The distinct patterns observed across niches demonstrate that adaptive microbiomes are not uniform, but finely tuned to plant physiology and environmental context, indicating tight plant–microbe co-evolution. This nuance is critical for understanding microbiome-mediated stress adaptation, which cannot be inferred solely from bulk soil or aboveground sampling.

Our results bridge mechanistic understanding of plant adaptation with applied significance. Serpentine soils can serve as a natural testbed to identify microbial taxa and plant genes that contribute to stress resilience. Understanding which plant loci drive microbial recruitment could allow future strategies to engineer crops that actively recruit beneficial microbes under nutrient-poor or metal-contaminated conditions. This moves beyond correlative microbiome studies: it suggests a pathway for predictive microbiome management, where plant genotype informs the assembly of adaptive microbial consortia.

## Online Methods

### Field sampling collection-strategy

Serpentine and nearby, genetically close non-serpentine control populations were collected in the Czech Republic and in Austria (**Fig. 1**). Populations N1 (49.734963, 15.174846, Vlastejovice) and S1 (49.683816, 15.133256, Borovsko) are located in the Czech Republic. Populations S2 (48.629930, 15.542567, Steinegg) and N2 (48.631497, 15.557237, Fuglau) are in Lower Austria, Austria and S3 (47.281679, 14.927647, Gulsen) and N3 (47.284170, 14.681940, Ingeringgraben) are in Styria, Austria. The selected populations were sampled in spring 2018 and fall 2018. For each population different tissues were sampled for their microbiomes: 1) leaf endophytes with closely associated epiphytes, 2) root endophytes with closely associated epiphytes, 3) rhizosphere and nearby 4) bulk soil. As microbial communities can change over time^80,81^, we sampled all tissues at each time-point. For this, leaves, roots, rhizosphere and bulk soil were collected in ten replicates. Additionally, soil and leaf material were collected for comprehensive ionomic profiling at each time point.

### Field sample collection – microbiome samples

All plant and soil material were collected using previously sterilised nitrile gloves and stored in 2 ml Eppendorf safe-lock tubes (Eppendorf) in RNAlater (SIGMA-ALDRICH) to minimise microbial gDNA degradation and community shifts. The sampled material was stored in a mobile refrigerator during transportation where the material was stored at -80 °C until use for gDNA extraction.

### Field sample collection – microbiome samples of transplant experiment

The transplant experiment is already published and described in^37,38^. In brief, to investigate adaptation mechanisms of serpentine populations of *A. arenosa* to serpentine soils Konečná et al grew each serpentine and its non-serpentine control population in a common garden set up in the experimental garden of the Charles University, Prague (Czech Republic) over the growing season of 2019. Each serpentine population was grown in its native serpentine soil and in the soil of its paired control non-serpentine population and vice versa. Specifically, serpentine populations (S1, S2, S3) were grown in their own serpentine soils (S1, S2, S3) as well as in the corresponding non-serpentine soils (N1, N2, N3). Likewise, non-serpentine populations (N1, N2, N3) were grown in their own soils and in the paired serpentine soils (S1, S2, S3). For example, population S1 was grown in its native serpentine S1 soil and in the paired non-serpentine N1 soil, and population N1 was grown in its native N1 soil and the paired serpentine S1 soil. In total 284 plants have been grown in total for the six populations, with 44-50 plants per population^37^.

To investigate the role of the plant microbiome in serpentine soil adaptation, leaf, root and bulk-soil material was collected for all populations grown in their natural and non-natural condition. All samples had been collected in September 2019 with five replicates per plant tissue, population and treatment. The collection and storing process was conducted as described above for microbiome samples.

### Field sample collection – ionomic profiling material

Soil and leaf material were collected for ionomic elemental profiling. The soil was collected into 15 ml conical tubes (Supplier Starlab (UK) Ltd). Leaves were collected into tea bags and stored in a box containing silica gel to reduce moisture. The material was stored at room temperature until preparation for elemental profiling.

### Ionomic material preparation

The leaf and soil material were prepared for ionomic profiling by air-drying the soil for a minimum of one week on the bench at room temperature and baking the leaves for three days at 60°C until the material was dry. The soil material was additionally sieved after baking to remove any debris. Leaf and soil samples underwent elemental profiling with Inductively Coupled Plasma Mass Spectrometry (ICP-MS) similar as described in^33,82^. Profiled elements were (names according to periodic table annotations): Ca, Mg, Mn, Ni, Cr, Rb, Co and Fe.

### Ion profile analyses

Ion concentration for elements of interest (namely Ca, Mg, Mn, Ni, Cr, Rb, Co and Fe) were visualised using custom scripts in R-4.4.1 with the ggplot2-3.5.1 package^83^. Statistic differences between serpentine and non-serpentine sites were tested using the stat_compare_means package of ggpubr-0.6.0^84^, specifying Wilcoxon rank-sum testing. The principal component analysis (PCA) was performed using the default R-4.4.1 prcomp function. The graph of variables was visualised using the fviz_pca_var function of factoextra-1.0.7^85^ with default settings and the ggplot2-3.5.1^83^ package in R-4.4.1.

### Map of populations

Geographic coordinates of the serpentine and non-serpentine populations were converted into spatial features using the sf-1.0 package in R-4.4.1, with WGS84 (EPSG:4326) as the reference coordinate system. For visualization, a base map of Austria and Czechia was obtained from the rnaturalearth-1.0.0 package at medium scale and used to display the location of populations^86^ with ggplot2-3.5.1^83^.

### Microbiome sample preparation

Samples were stored in RNAlater^82,87–90^. Roots and leaves were transferred into a fresh 2 ml tube and washed with 2 ml 0.1 M potassium phosphate pH 8.0 buffer with shaking at 200 rpm for 30 min. Afterwards the root was transferred into a fresh 2 ml tube and supernatant was retained as the rhizosphere sample. This step was repeated three times, with fresh buffer each time. Afterwards the sample was vortexed twice for 30 seconds with fresh 2 ml 0.1 M potassium phosphate pH 8.0 buffer and the root was transferred into a fresh 2 ml tube. The supernatant was added to the previously collected supernatant. The combined supernatant was centrifuged for 10 min with 15,000 rcf and room temperature. After removing the supernatant, the pellet was kept as a rhizosphere sample and stored at -80°C until further use. Leaf samples were washed with 2 ml 0.1 M potassium phosphate pH 8.0 buffer by vortexing twice for 10 sec. This step was repeated three times. The supernatant was decanted, and leaves transferred into a fresh 2 ml tube and stored at -80°C until further use.

### Microbiome gDNA extraction of field and transplant experiment samples

The gDNA extraction for the field and common garden experiment was performed as described by Bollmann-Giolai *et al.*^51^ using 250 mg soil, before gDNA was purified in a 96-well format from 140 µl supernatant using magnetic beads (Sera-Mag Carboxylate-Modified Magnetic Particles (Hydrophophilic), GE Healthcare Life Sciences). The cleaned gDNA was transferred to a fresh 96 well plate and stored at -20 °C until further use.

### Wet lab – Amplicon bacterial and fungal library construction

Amplicon data was analysed as previously described^23,51^. In brief: The bacterial variable (V4) region of the 16S rRNA gene and the fungal ITS1 region of the Internal Transcribed Spacer were targeted for amplicon library construction using 16S 515 forward, 16S 806 reverse, ITS 1 forward and ITS2 reverse primers adapted from Walters et al ^91^ (**Extended Data File 10**). Amplicon library construction was performed using a two-step PCR protocol as previously described^23,51^: In a first PCR the 16S V4 and ITS regions were targeted using gene specific primers, followed by custom dual indexing in a barcoding PCR (**Extended Data File 10**). For the bacterial 16S V4 PCRs PNAs (peptide nucleic acid PCR clamps) were included targeting plant mitochondria (mPNA) and chloroplast regions (cPNA)^92^ in the first PCR to minimise plant gene amplification. Library construction was conducted as previously described^23,51^. Briefly, the first PCR step was performed with three technical replicates using 3 ng gDNA. Samples were then pooled and subjected to a 0.7x magnetic bead clean-up (MAGBIO) and eluted in 10 µl TE buffer. The second PCR was conducted using the program for bacterial and fungal libraries previously described^23,51^.

The cleaned amplicon libraries were quantified using the Qubit 2.0 fluorometer (Thermo Fisher Scientific) with dsDNA HS Assay Kit reagents (Thermo Fisher Scientific) and the size of the amplicons was controlled on the GX Touch using the 3K kit (X-Mark DNA LabChip, HT DNA NGS 3K Reagent Kit, Perkin Elmer LAS (UK) LTD). Amplicons were pooled equimolarly to 1.5 nM for bacterial and 2.0 nM for fungal libraries according to the molarity. A final 0.7 x magnetic bead clean-up (HighPrep^TM^ PCR Clean-up System, MAGBIO) was performed and the final library pools were eluted in 50 µl EB buffer.

### Sequencing on NovaSeq SP flowcell

The libraries were equimolarly pooled for NovaSeq sequencing. Prior to submission the final library pools were concentrated with a 0.7x magnetic bead clean-up and eluted in 50 µl EB buffer. The libraries were sequenced on one SP flow cell on the NovaSeq with a 30 % PhiX spike-in.

### Amplicon sequence variant taxonomy assignment and filtering

Amplicon data analysis was conducted as previously described^23,51^ with minor alterations: The Illumina binary base call files (bcl) were demultiplexed using bcl2fastq version 2-2.20.0.422 with the settings - -barcode-mismatches 1 --fastq-compression-level 9 into individual fastq.gz files. The paired-end reads were trimmed for primers, sequencing adapters and linker sequences using cutadapt-1.9.1^93^ with the settings -n 4 --minimum-length=50. Following cutadapt fastp-0.20.0^94^ was run on 16S data with disabled length filter and trim_poly_g to trim polyG read tails. Afterwards both 16S and ITS data was quality controlled, trimmed and filtered using R-3.6.3 and DADA2 version 1.14.1 according to the workflow described in version 2^95^. The truncation length for forward reads was set to 210 bp and the truncation length for the reverse reads to 210 bp for both 16S and ITS. The truncation length was based on read quality. For 16S and ITS libraries we used the following parameters: maxN=0, maxEE=c(2, 2) and truncQ=2 and a minimum length of 50 bp. The filtering parameter maxEE chosen allows for a maximum of 2 expected errors per read, the filtering parameter maxN chosen does not allow any Ns. In addition to filtering using maxEE, maxN and read trimming, a quality score (truncQ) was set that truncates the read at the first nucleotide with a quality score of 2 (set value). After these filtering steps, error rates were estimated as described in^95^ and the data was denoised with standard inference settings^95^. Forward and reverse reads were merged with default settings: if the minimum of 12 identical bases for overlap was not met between forward and reverse reads, the sample would be omitted. Following the merging of paired reads, a sequence table was constructed with amplicon sequence variants (ASVs). As an additional, subsequent filtering step, chimaeras were removed using standard settings^95^. The Silva (silva_nr99_v138.1.train.set.fa)^96^ and UNITE (sh_general_release_dynamic_all_04.04.2024.fasta)^97^ databases were then used to taxonomically classify bacterial and fungal reads respectively. Reads that did not match either the bacterial or fungal database were removed.

Amplicon sequence variants (ASVs) were used to classify both bacterial and fungal reads. A minimum read number after filtering of 1000 reads per sample and a minimum of three samples per dataset was set as cut-off. We further applied a hard-filter removing all ASVs which were not supported by at least 10 reads per sample and a total relative abundance of 0.1 %. Filtering reduced the number of reads in our 16S libraries from an initial average of 177,387 raw reads to 42,854 reads per library after filtering and assignment to ASVs. For ITS libraries, we initially obtained an average of 77,998 raw reads, which was reduced to 65,666 reads per library following taxonomic assignment. After quality controlling, trimming, taxon assignments and filtering, which were all carried out on the Norwich high-performance computing cluster, we continued with analyses on a desktop machine in R-4.4.1.

### Alpha- and beta-diversity comparisons

Alpha-diversity (Shannon and Observed measures) was calculated on pre-normalised data using the phyloseq-1.48.0 package^98^. Statistical significance of diversity differences was assessed with Kruskal-Wallis rank sum tests and pairwise Wilcoxon rank sum tests for non-normally distributed data, or with ANOVA followed by Tukey’s HSD for normally distributed data, using default functions in R-4.4.1.

For beta-diversity analysis, read counts were transformed using the transform_sample_counts function of phyloseq-1.48.0^98^. Bray-Curtis dissimilarities were calculated with the vegan-2.6 package in R-4.4.1 and community-level differences were tested using ANOSIM and PERMANOVA (adonis function) with the vegan-2.6 package^99^. To visually assess differences between niches we visualised samples using RDA plots in R-4.4.1 and custom ggplot2-3.5.1^83^.

### Relative abundance plots across niches and conditions

To show relative abundance plots of phyla across plant niches, we used custom R-4.4.1 scrips. We agglomerated taxa to phyla using the phyloseq-1.48.0^98^ tax_glom function and visualised relative abundances per phylum as percent-of-all ASVs using ggplot2-3.5.1^83^.

### Calculating sharedness of ASVs

The sharedness of microbial ASVs between serpentine or non-serpentine populations were calculated using custom R-4.4.1 scripts. ASVs that were found in all each of three serpentine or non-serpentine populations were assigned to the conserved group. ASVs present in two populations were assigned to the shared group. ASVs present in only one population were annotated as unique. The data were visualised using ggplot2-3.5.1^83^ in R-4.4.1.

### Differential abundance testing of ASVs

Differential abundance (i.e., enrichment) testing between serpentine and non-serpentine conditions for each niche was performed using the default microbiomeMarker-1.10.0^55^ settings with its run_limma_voom and marker_table functions in R-4.4.1. Enrichment tables were expanded by connecting ASVs with their taxonomy using custom R-4.4.1 scripts and visualised the data either as a volcano-plot or scatterplots comparing adjusted p-values using custom ggplot2-3.5.1^83^. The comparison between field and transplant enrichment (i.e., log2 fold-change values obtained by the run_limma_voom) function was visualised using ggplot2-3.5.1^83^. Spearman’s correlation coefficients were calculated using the stat_cor function of ggpubr-0.6.0^84^ in R-4.4.1.

### Bray-Curtis comparison of microbiomes

The Bray-Curtis distance of microbiomes from transplanted populations was calculated using the vegdist function of the vegan-2.6 package^99^ in R-4.4.1. For each population and niche relative abundance values of genera detected under the serpentine as well as non-serpentine conditions were combined to an abundance matrix. The abundance matrix was analysed using the vegdist function. The obtained results were visualised using custom ggplot2-3.5.1 scripts.

### Estimating microbial or genotype variances

To determine genotype profiles of serpentine and non-serpentine populations we compared k-mers of *A. arenosa* genomes previously collected from each population using kWIP-0.2.0^100^ default settings. This produced a distance matrix, which was analysed using the default cmdscale function in R-4.4.1. The output of the Classic Multi-dimensional Scaling (MDS) analysis was visualised using ggplot2-3.5.1^83^. This resulted in distinct separation of genotypes by population on the MDS plot.

To obtain a single environmental variable describing serpentine versus non-serpentine stress across all tested elements, the previously calculated PC1 values of populations were used. These values captured variation in calcium/magnesium stress and heavy metal composition.

To estimate environmental and genotype contributions to bacterial and fungal community assembly, all ASVs that were statistically significantly enriched in the field experiment under either serpentine or non-serpentine conditions were retained. Based on the total relative abundance of each ASV, generalized linear mixed models (GLMMs) were fitted using the glmer function of the lme4-1.1^101^ package in R-4.4.1. To disseminate contributions of the genotype or the environment to microbial abundance three models were tested. (1) A full model, in which microbial abundance was explained by both population identity and condition, with time included as a random effect. (2) A population-only model, in which microbial abundance was explained by population identity and time as a random effect. (3) An environment-only model, in which microbial abundance was explained by condition and time as a random effect. All models were fitted using a Gamma distribution with a log link function. Optimization was performed using a maximum number of iterations of 1e5. For each fitted model with a statistically significant contribution of genotype or environment to microbiome assembly, conditional R^2^ values were calculated using r2 function of the performance-0.12.4^102^ package in R-4.4.1. Model coefficients were extracted and R^2^ values visualised using custom ggplot2-3.5.1 scripts.

## Acknowledgements

We thank Iain Macaulay for helpful discussion about the newest genomic methodologies, Alba Pacheco-Moreno for discussions about plant-microbe interactions and David E. Salt for his support with elemental profiling.

## Data Availability Statement

The sequencing data can be accessed at the NCBI BioProject database (BioProject ID: PRJEB71077).

## Additional Files

**Extended Data File 1**: Statistical analysis of ion compositions in serpentine and non-serpentine soils.

**Extended Data File 2**: Word document containing supplementary figures, alpha- and beta-diversity analyses.

**Extended Data File 3**: Percentages of bacterial and fungal phyla detected across seasons, populations, and niches.

**Extended Data File 4**: Percentages of conserved, shared, and unique fungal and bacterial ASVs across populations.

**Extended Data File 5**: Table displaying percentages of bacterial and fungal ASVs enriched in serpentine and non-serpentine soils.

**Extended Data File 6**: Results from field enrichment testing.

**Extended Data File 7**: ASVs per genus from the field enrichment experiment.

**Extended Data File 8**: Comparison of transplant and field enrichment testing results.

**Extended Data File 9**: Overlap of enrichment testing for serpentine-enriched bacterial and fungal ASVs between field and transplant.

**Extended Data File 10**: Excel file containing sequencing adapters, barcodes, and primers.

