## Extended data file 2 for "*Arabidopsis arenosa* influences its microbiome as a serpentine soil adaptation strategy"

**Appendix file 1**

**Alpha-diversity analyses**

Alpha-diversity analyses of 16S data showed that neither Observed richness nor Shannon diversity differed significantly across serpentine and non-serpentine conditions in spring (Observed p = 0.877; Shannon p = 0.994) or fall 2018 (Observed p = 0.342; Shannon p = 0.226) indicating that 16S microbial community diversity was consistent across conditions in both seasons. For ITS data Shapiro-Wilk tests indicated that both measures were non-normally distributed in spring (Observed p = 0.039; Shannon p = 8.48 x10^-5^) and fall (Observed p = 0.028; Shannon p = 7.1 x10^-4^), so non-parametric tests were applied. Kruskal-Wallis tests showed no significant differences across conditions for either spring (Observed p = 0.728; Shannon p = 0.373) or fall (Observed p = 0.909; Shannon p = 0.971), indicating that fungal community richness and overall diversity were consistent across conditions in both seasons.

For 16S spring populations, Shapiro-Wilk tests indicated non-normality for Observed richness (p = 2.57 x10^-7^) but approximate normality for Shannon diversity (p = 0.388); neither Kruskal-Wallis (Observed p = 0.736) nor ANOVA (Shannon p = 0.831) detected significant differences between populations. For 16S fall populations, both diversity measures were non-normal (Observed p = 1.03 x10^-6^; Shannon p = 0.008), and Kruskal-Wallis tests showed a significant difference in Observed richness among populations (p = 0.048) but not in Shannon diversity (p = 0.078); however, pairwise Wilcoxon comparisons revealed no significant differences between any population pairs. For ITS data, Shapiro-Wilk tests indicated non-normality for both spring (Observed p = 0.039; Shannon p = 8.48 x10^-5^) and fall (Observed p = 0.028; Shannon p = 7.10 x10^-4^) populations, and Kruskal-Wallis tests showed no significant differences in either Observed richness or Shannon diversity across populations in spring (Observed p = 0.841; Shannon p = 0.212) or fall (Observed p = 0.104; Shannon p = 0.068). Overall, these results indicate that microbial community alpha-diversity remained broadly consistent across populations, with only minor differences in 16S richness observed in fall.

Alpha-diversity of microbial communities was assessed across different plant-associated niches (bulk soil, leaf, rhizosphere, and root). Shapiro-Wilk tests indicated that both Observed richness and Shannon diversity were non-normally distributed for ITS spring (Observed p = 0.039; Shannon p = 8.48 x10^-5^), ITS fall (Observed p = 0.028; Shannon p = 7.1 x10^-4^), 16S spring (Observed p = 2.57 x10^-4^) and 16S fall (Observed p = 1.03 x10^-6^; Shannon p = 0.008), while Shannon diversity in 16S spring was normal (p = 0.388). Kruskal-Wallis tests revealed significant differences in both Observed richness and Shannon diversity across niches for all datasets (ITS fall: Observed p = 0.00077, Shannon p = 0.00091; ITS spring: Observed p = 2.95 x10^-10^, Shannon p = 8.52 x10^-9^; 16S fall: Observed p = 4.45 x10^-8^, Shannon p = 2.33 x10^-6^; 16S spring: Observed p = 0.00013), except for 16S spring Shannon, which was tested by ANOVA (p = 0.064) and showed no significant differences. Pairwise Wilcoxon tests indicated that leaf samples consistently harboured significantly lower diversity compared to other niches in most datasets, while differences between rhizosphere and root were more variable depending on season and dataset. Overall, these results demonstrate that microbial alpha-diversity varies strongly across plant-associated niches, with the leaf generally supporting lower richness and diversity than belowground compartments.

Altogether this indicates that 16S and ITS community diversity was generally consistent across serpentine and non-serpentine conditions in both spring and fall 2018, without significant differences detected in Observed richness or Shannon diversity. Across populations, 16S communities showed only minor differences in richness during fall, while ITS diversity remained consistent between populations in both seasons. In contrast, microbial alpha-diversity varied strongly across plant-associated niches, with both 16S and ITS datasets showing significantly lower richness and diversity in leaf samples compared to belowground compartments, while rhizosphere and root differences were more variable depending on season and dataset. Overall, these results indicate that niche exerts a strong influence on microbial alpha-diversity than condition or population.

**Beta-diversity analyses**

Beta-diversity analyses showed that microbial community composition differed depending on population, condition, and plant-associated niche, with patterns varying by dataset and season. For 16S spring, adonis detected significant effects of population (p = 0.001) and a weaker effect of condition (p = 0.046), with ANOSIM again showing strong niche separation (R = 0.546, p < 0.001) and permutest confirming significant differences across niches (p < 0.001) but not populations or condition. 16S fall, adonis tests indicated significant effects of both population and condition on community composition (p = 0.001), and ANOSIM revealed strong separation across niches (R = 0.573, p < 0.001), supported by significant differences in permutest for material (p < 0.001) but not for population or condition. In ITS data showed similar trends: in spring, adonis revealed significant effects of population (p = 0.001) and condition (p = 0.009), with ANOSIM confirming strong niche differences (R = 0.238, p < 0.001) and permutest indicating significant material effects (p < 0.001). For ITS fall, adonis identified condition effects (p = 0.003), while ANOSIM indicated weaker separation for population (R = 0.172, p < 0.001) and material (R = 0.247, p < 0.001), and permutest confirmed significant differences for population (p = 0.024) and material (p < 0.001). Overall, these results indicate that plant-associated niche consistently exerts the strongest influence on microbial community composition, while population and condition have smaller but occasionally significant effects depending on season and dataset.

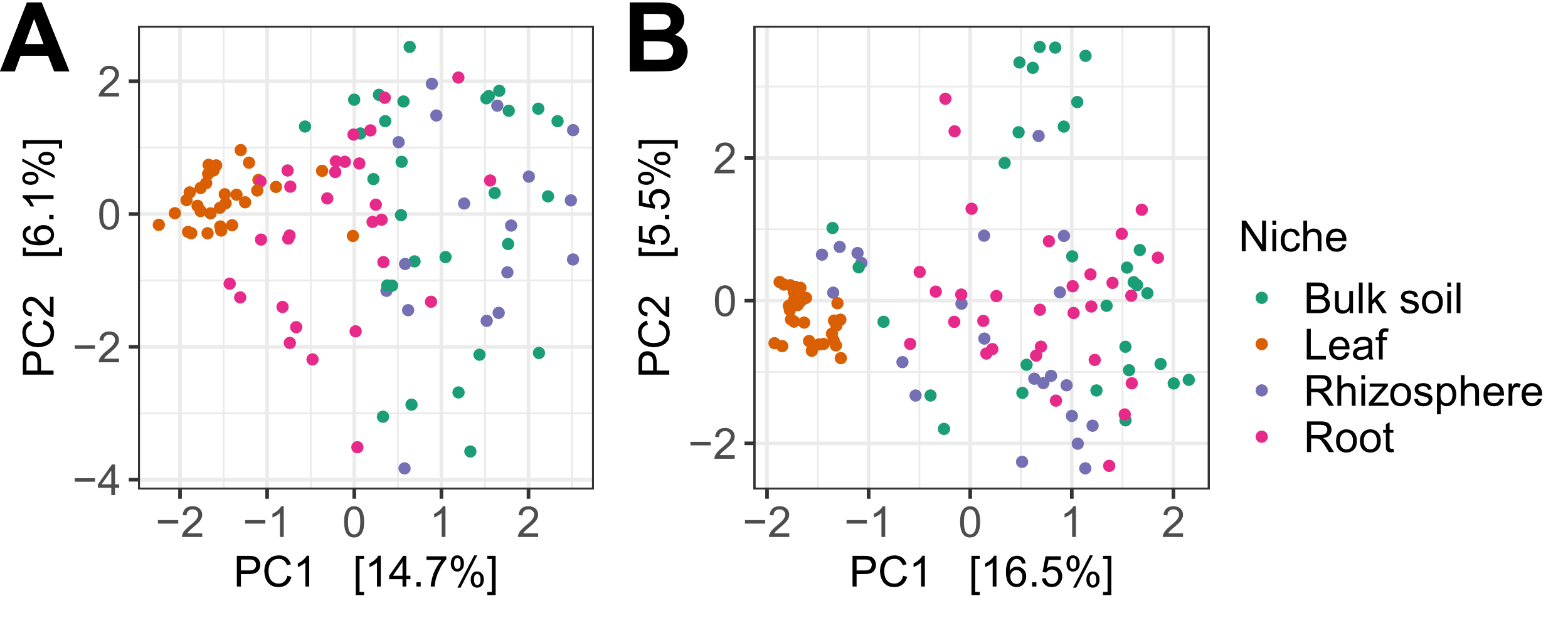

**Additional Figure 1** RDA of **(A)** bacterial and **(B)** fungal communities in bulk soil, rhizosphere, root, and leaf niches in spring 2018.

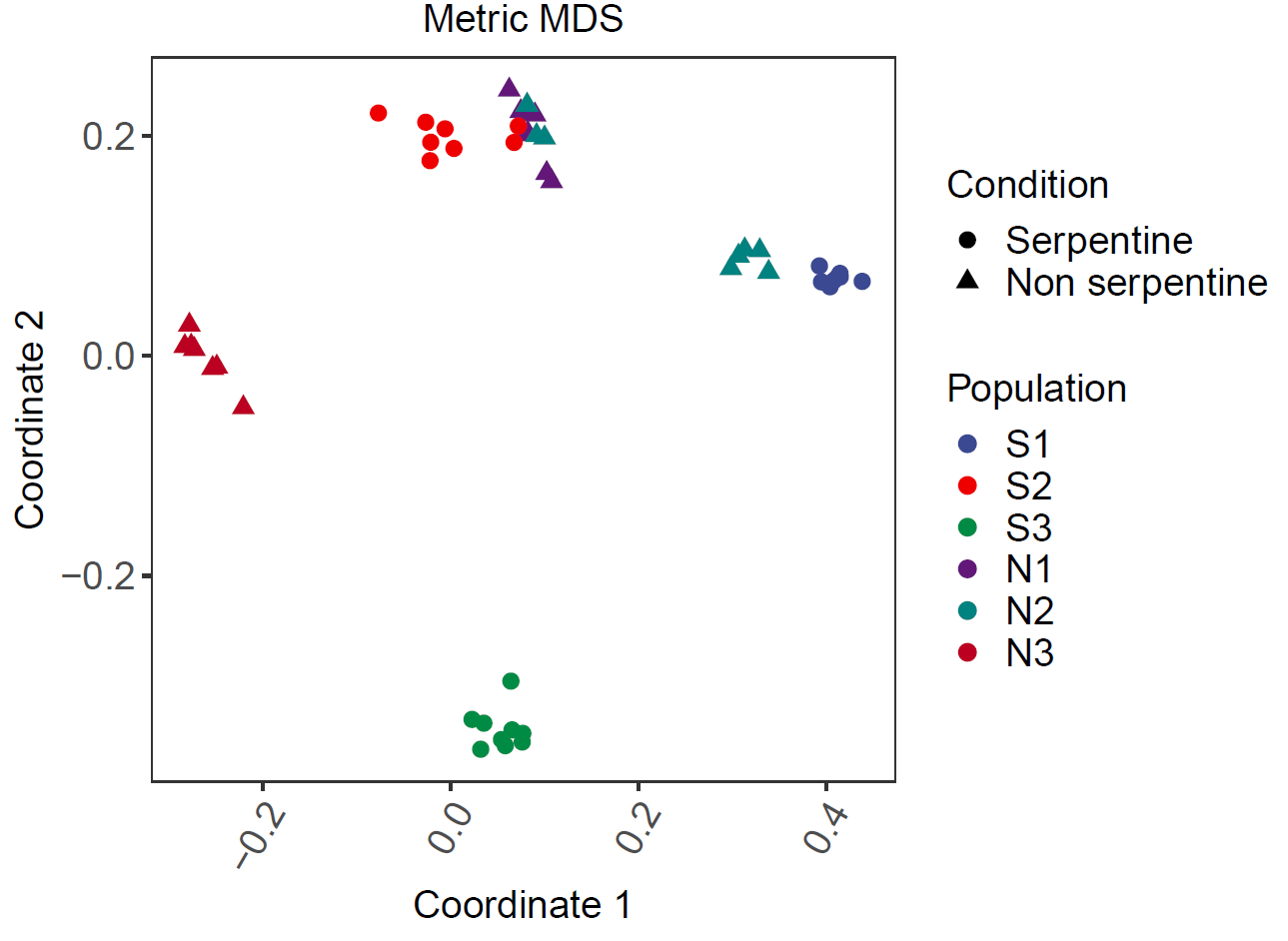

**Additional Figure 2 Genetic relationships among individuals estimated from sequencing reads using kWIP.** Multidimensional scaling (MDS) of genome-wide k-mer-based genetic similarity shows that most populations form distinct groups. Within each population, individuals are more closely related to one another than to individuals from other populations.
